# Inchworm stepping of Myc-Max heterodimer protein diffusion along DNA

**DOI:** 10.1101/2020.06.17.156398

**Authors:** Liqiang Dai, Jin Yu

## Abstract

Oncogenic protein Myc serves as a transcription factor to control cell metabolisms. Myc dimerizes via leucine zipper with its associated partner protein Max to form a heterodimer structure, which then binds target DNA sequences to regulate gene transcription. The regulation depends on by Myc-Max binding to DNA and searching for target sequences via diffusional motions along DNA. Here, we conduct structure-based molecular dynamics (MD) simulations to investigate the diffusion dynamics of the Myc-Max heterodimer along DNA. We found that the heterodimer protein slides on the DNA in a rotation-uncoupled manner in coarse-grained simulations, as its two helical DNA binding basic regions (BRs) alternate between open and closed conformations via inchworm stepping motions. In such motions, the two BRs of the heterodimer step across the DNA strand one by one, with step sizes up about half of a DNA helical pitch length. Atomic MD simulations of the Myc-Max heterodimer in complex with DNA have also been conducted. Hydrogen bond interactions reveal between the two BRs and two complementary DNA strands, respectively. In the non-specific DNA binding, the BR shows an onset of stepping on one association DNA strand and dissociating from the complementary strand. Overall, our simulation studies suggest that the inchworm stepping motions of the Myc-Max heterodimer can be achieved during the protein diffusion along DNA.

## 1. Introduction

Gene expression relies essentially on transcription factors (TFs) search on DNA to locate sequence motifs for subsequent regulation protein association and assembly. During the search process, the TF protein often binds to non-specific DNA and then approaches to a specific DNA binding site via diffusion along the DNA [1, 2]. A facilitated diffusion model had been suggested in which TFs alternate between 3-D cellular space diffusion and 1-D diffusion along DNA in order to achieve efficient target search and binding [1, 3-7]. In 1-D diffusion, TF can track, slide, and hop along DNA. It can also jump for intersegmental DNA transfer.

Although phenomenological models of the facilitated diffusion have been established [1, 3-7], molecular details of the TF diffusion remain largely unknown. Single molecule experiments could detect TF diffusional motions on the DNA [8-10], yet it is highly challenging to capture the movements at base-pair (bp) spatial resolution or sub-millisecond time resolution. In comparison, motor proteins can be slowed down in a controllable manner (e.g. by reducing ATP concentration) to allow high-resolution detection. For example, real-time single-molecule measurements demonstrated hand-over-hand stepping motions of kinesin or myosin motors along microtubes or actin filaments [11-14]. Single bp movements of nucleic acid motors such as RNAP polymerase [15] or DNA helicase along DNA track have also been identified [16, 17].

On the other hand, structure-based simulations allow for molecular dynamics (MD) characterization of key structure components of the systems. Atomic MD simulations contain finest structural dynamics details but are often limited by simulation time scale, which can hardly surpass several microseconds for a TF protein-DNA system of a regular size [18-20]. Coarse-graining (CG) techniques, however, provide ways to extend the simulation time scale via maintaining essential protein-DNA electrostatic interactions to support the TF diffusion [21-24]. In current studies implementing the CG simulations using CafeMol [25], each protein amino acid is represented by one bead, and each DNA nucleotide is modeled by three beads [26]. Similar simulation studies have demonstrated rotation-coupled or uncoupled sliding motions of TFs along DNA [22, 27, 28].

Here, we employed mainly the protein-DNA structure-based CG simulations to investigate diffusional motions of a heterodimer TF Myc-Max on the DNA. Myc plays an important role in cell proliferation, differentiation, and apoptosis. Deregulated level of Myc leads to abnormal cell growth or cancer [29-31]. Myc cannot form a homodimer under physiological condition, instead, it forms a heterodimer with the Myc-associated-factor X (Max) for most of known functions [32-34]. The Myc-Max heterodimers can specifically bind to cognate DNA sequences (e.g. 5’-CACGTG-3’) termed the enhancer box or E-box [32]. In regular conditions, Myc-Max binds mostly to the E-box and consensus sequences populated in active promoter regions [35].

The Myc family members contain a N-terminal transactivation domain, a middle segment rich in proline, glutamic acid, threonine, and proline residues (PEST), and a C-terminus basic region/helix–loop–helix/leucine zipper (bHLHLZ) domain [36, 37]. The N-term and PEST domain of Myc are largely disordered. The well-folded molecular structure of the bHLHLZ domain is presented in **Fig 1A** [38]. The Myc/Max monomer structure thus contains the DNA-bounded basic region (BR), the helix-loop-helix (HLH) motif, and the leucine zipper (LZ) domain that severs as an anchor to stabilize the dimer structure [38, 39]. The presented structure appears to be in a closed state. An open conformation of homodimer Max-Max has also been suggested [40].

**Fig 1.**
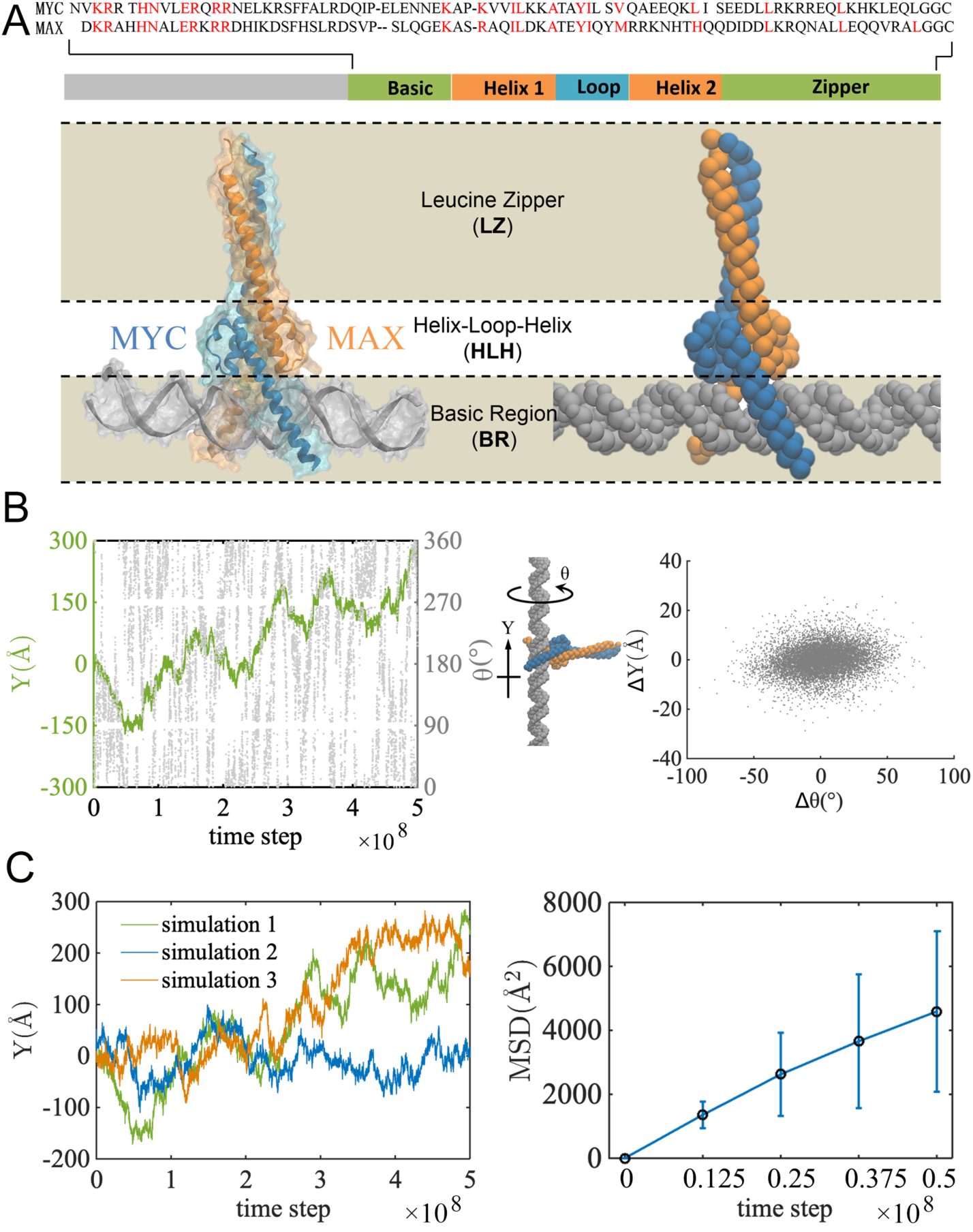
Coarse-grained (CG) simulations of the Myc-Max heterodimer diffusing along DNA. (A) The Myc and Max protein sequence alignment is shown and the conserved residues are highlighted in red *(top)*. The molecular view of the heterodimer in complex with DNA (pdb: 1NKP) [38] is shown *(bottom left)*, with Myc, Max, and DNA colored in blue, orange, and gray, respectively. The CG structure of the complex is also shown *(bottom right)*. (B) The center of mass (COM) positioning of Myc-Max BRs mapped on the DNA. The DNA long axis is aligned to the Y-axis. The position of the protein along the DNA is shown by green line, and the protein rotational degree around the DNA is shown in gray dots (*left*). For every 1000 timesteps, 2D mapping of the longitudinal distance ΔY and the rotational degree Δθ is shown (*right*). (C) The diffusion of the protein along DNA. The COM of the BRs along the DNA for three independent runs are shown *(left)*. The mean square displacements (MSDs) of the protein COM vs the time is also presented for the simulations *(right)*.

Computational studies have been conducted mostly toward Myc-Max inhibition for drug discovery approaches [41-43]. The dimerization specificity of c-Myc had been modeled [44]. Modeling also shows different promoter binding affinities accounting for Myc gene regulation specificity [45]. In this work, we focus on characterizing diffusional motions of the Myc-Max heterodimer along nonspecific DNA. To do that, we conducted the structure-based CG simulations, in which the protein-DNA electrostatic interactions are largely maintained. We found that the Myc-Max heterodimer transits between two dominant conformations (closed and open), and two DNA binding BRs alternately step across the DNA strand in rotation-uncoupled inchworm motions [46, 47]. Additional atomistic MD simulations up to one microsecond each have also been conducted for Myc-Max in complex with specific and non-specific DNA. Initiation of stepping and DNA strand dissociation of Myc show in the non-specific DNA binding system.

## 2. Results

Here we present the simulations that reveal the inchworm stepping motions of Myc-Max during 1-D diffusion along DNA. We first examine whether protein translational motions are coupled with DNA helical tracking or rotational motions. Then we focus on analyzing conformational changes of the Myc-Max heterodimer, particularly on the opening and closing between the two BRs of the heterodimer during the diffusion. Based on these examinations, we then present an inchworm stepping model of the Myc-Max diffusion, with stochastic BR swapping events described as well. We additionally present all-atom MD simulations of the Myc-Max heterodimer on the specific and non-specific DNA, probing whether there are clues to protein diffusional motions in the non-specific DNA binding.

### 2.1 The rotation uncoupled sliding of the Myc-Max heterodimer along DNA

We conducted CG simulations to investigate the Myc-Max diffusion along DNA. The CG protein was built by using the Go” odel [48], with each amino acid represented by one bead. The CG DNA is built by 3SPN2C model [26], with each nucleotide represented by 3 beads, corresponding to sugar, base and phosphate, respectively (**Fig 1A** *bottom*). The simulation was conducted for 5×10^8^ time steps (∼10 μs [25] at least) under a physiological ionic condition (150 mM). The protein is initially placed 25Å above DNA, then quickly binds to the DNA and starts diffusing along the DNA in the simulation (see **Supplementary Information** or **SI Fig 1**). The positioning of the center of mass (COM) of the protein BRs is measured (**Fig 1B** *left*). The protein moves and rotates occasionally around the DNA during sliding in both directions (i.e., +Y toward left and -Y right). Although the Myc-Max heterodimer could rotate around the DNA from time to time, the correlation between the protein longitudinal sliding on the DNA (ΔY) and the rotational (Δθ) per 1000 timesteps appears low (correlation coefficient=0.16; **Fig 1B** *right*). Hence, the sliding of Myc-Max is rotationally uncoupled. To further investigate the diffusion rate of the protein along DNA, two additional CG simulations were conducted (**Fig 1C** *left*). The mean square displacements (MSDs) of the COM of protein BRs along DNA are shown (**Fig1C** *right*), square displacements (MSDs) of the COM of protein BRs along DNA are shown (**Fig 1C** *right*), with a diffusion coefficient fitted at to ∼ 4.6×10^−5^ Å^2^/step or 23 nm^2^/μs. Note that the diffusion coefficient can be overestimated due to lack of detailed interactions between protein and DNA in the CG simulations.

How fast Myc-Max diffuses also depends on the ionic condition. Under a low ionic concentration, due to weak charge screening, the protein is bound tightly to the DNA. Interestingly, we notice that the dimer may dissociate into two monomers, which slide independently along DNA in the CG simulations (see **SI Fig 2A**). Previous studies have shown that Myc and Max monomers can bind DNA first and dimerize on the DNA due to a kinetic preference [49], while the dimer dissociation captured here appears to be an inverse process. Additionally, under a high ionic concentration (200 mM), with strong charge screening on the protein-DNA association, the protein dissociates from DNA (**SI Fig 2B**).

### 2.2 Dominant conformational changes of Myc-Max during diffusion along DNA

The sliding of the Myc-Max heterodimer is however coupled closely to significant conformational changes between the two BRs. During the sliding, the two BRs undergo scissor-like motions to open and close so that to step along DNA. We measure the distance between the two BR COMs of Myc and Max on the DNA as dY (see **Fig 2A** *left*). Within a long period of time, a certain ‘left-right’ positioning of the Myc-Max BRs is maintained (e.g. dY>0 for Myc/Max on the left/right lasts for 7.5×10^5^ timesteps).

**Fig 2.**
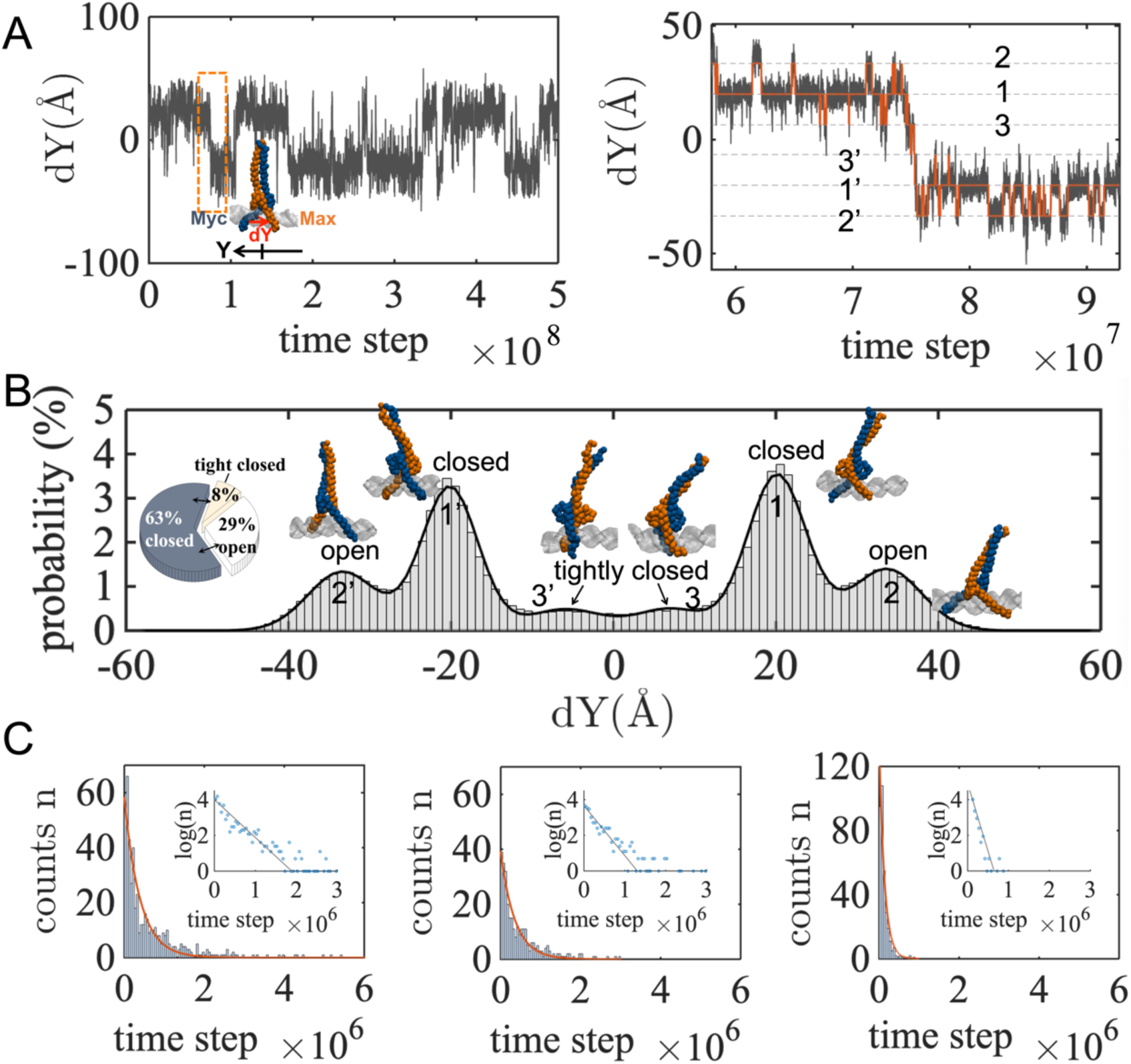
Various inter-BR conformations of Myc-Max during diffusion along DNA. (A) The distance vector dY between the COMs of the two BR domains of Myc and Max chain is mapped along the DNA during the CG simulation 1 *(left)*. The zoom-in view of the trajectory within an orange rectangle is shown (*right*). Six states are labeled, three for dY>0 (1, 2, and 3) and three for dY<0 (1’, 2’, and 3’). (B) The histogram of the distance vector dY between the Myc and the Max BRs is shown, with a representative conformation for each peak illustrated. (C) The survival conformational counts over time steps for the open, closed, and tightly closed states measured from the simulation 1, the average survival or duration time for each state is obtained.

For a complete CG simulation 1, we find that dY(t) is not simply a two-state series, but is composed of six states (**Fig 2A** *right*; with labels 1-2-3 for dY>0 and 1’-2’-3’ for dY<0). The histogram of dY from the full simulation is shown (**Fig 2B**). There are four significant peaks and two insignificant but visible ones in the histogram, referring to three inter-BR conformations (closed for 1 and 1’, open for 2 and 2’, and tightly closed as 3 and 3’): (1) The closed state (|dY|∼20±3.5Å) resembles the crystal structure [38], with the two BRs bound on the two sides of a same DNA groove; (2) the open state (|dY|∼33±4.2Å) is reached as the leading BR moves across the DNA backbone and bind to the next DNA groove, starting from the closed state; (3) the tightly closed state (|dY|∼6.5±4.4Å) is obtained as the lagging BR moves further close to the leading BR, also starting from the closed state. The closed state accounts for ∼ 63% of the overall population, with a duration time ∼ 4.1×10^5^ steps (see **Fig 2C**); the open state accounts ∼ 28%, with a duration time ∼2.5×10^5^ steps; and the tightly closed state accounts <10%, with a duration time 0.9×10^5^ steps.

### 2.3 An inchworm stepping model of the Myc-Max heterodimer diffusion along DNA with stochastic left-right repositioning or reversal

Correspondingly, we suggest that the Myc-Max heterodimer diffusion along DNA follows an inchworm model (see **Fig 3A)**, moving either backward or forward (see **SI Movie S1**/**S2**). Starting from the highly populated closed conformation (state 1), the leading BR moves forward first across the DNA backbone to the next DNA groove, at a step size of ∼13 Å, so that Myc-Max transits to the open conformation (state 2); then the lagging BR follows to recover the protein back to the closed state. Such BR stepping motions represent typically the inchworm type of domain/subdomain motions [46, 47].

**Fig 3.**
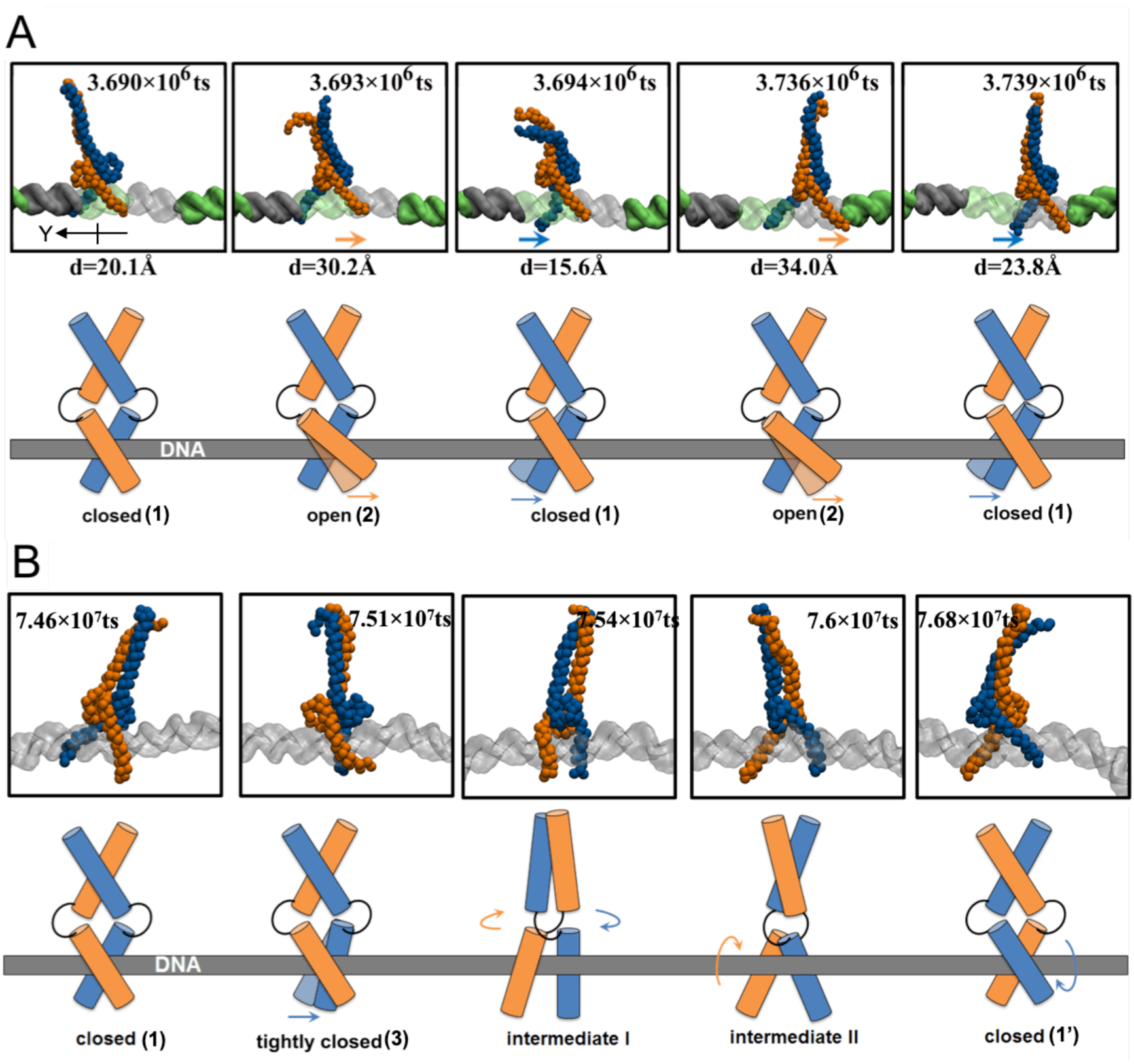
An inchworm stepping model of Myc-Max diffusion on the DNA. (A) The two BRs of the Myc-Max heterodimer step along DNA. The representative snapshots are taken from the CG simulation 1 *(top)*, with distances between the Max and Myc BRs labeled. The Myc chain is colored in blue and the Max in orange. The DNA is shown in the surface representation, colored in gray and green alternately for every 10 bp. The schematics of the Myc-Max inchworm stepping with alternating closed and open conformations are also shown *(bottom)*. (B) The Myc-Max *left-right* positioning reversal or BR swapping captured in the simulation (*top*) and illustrated in schematics (*bottom*), with two closed and one tightly closed state (1, 1’ and 3 in Fig 2) and two intermediate states (I and II) presented.

Occasionally, starting from the closed state, the lagging BR can also move forward first. In such a case, the heterodimer transits to the tightly closed state (state 3), which is of low population and shortly lived. Furthermore, rather than transiting back to the stabilized closed state, the tightly closed state also allows the BR ‘swapping’ so that the left-right positioning between the two BRs exchanges or reverses. Such BR swapping events (e.g. 134 caught in the 5 × 10^8^ simulation timesteps; see **SI Movie S3**) stochastically intervene the regular inchworm stepping. Followed by the BR swapping, the other closed state (1’) with an opposite left-right positioning is reached, via protein self-rotation and two intermediate states (I and II, see **Fig 3B** bottom). The protein rotation seems to be induced by a torque upon tightening of the two BRs (i.e. from state 1 to 3). As the rotation leads to the intermediate state I, the BR from right moves toward left, but can hardly insert into the DNA groove as it is against the direction of the DNA strand; it then flips across to the other side of the DNA helix, so that two BRs stay on the same side of the DNA, reaching the intermediate states II. Finally, the torque can be released after the flipping of the other BR across the DNA. In this way, the two BRs of Myc and Max reverse their left-right positioning (from state 1 to 1’).

It is noted that overall the probabilities for Myc and Max becoming a leading chain in the diffusion keep the same (see **SI Fig 3**). It is equally likely for Myc-Max to move forward and backward, and it is also equivalent to place Myc and Max left or right in the heterodimer on the double-stranded (ds)DNA without structure or sequence bias.

### 2.4 Binding and onset of diffusion of Myc-Max on DNA in atomistic MD simulations

Although the CG simulations capture essential protein-DNA electrostatic interactions, the hydrogen bond (HB) interactions formed at protein-DNA interface are missing. To include those detailed interactions and probe the corresponding dynamics, we conducted respective 1-µs atomistic MD simulations for Myc-Max binding onto specific (E-box) and nonspecific (ploy-AT) DNA sequences. The initial and final structures of the Myc-Max heterodimer bound on the DNA are presented (see **Fig 4A**). The long LZ domain deviates most significantly from that in the initial structure. The deviations of the full heterodimer and non-LZ domain are both larger than that on the specific DNA (**Fig 4B**). The domain also fluctuates most, and the fluctuations of the protein on the specific DNA is obviously smaller than that on the nonspecific DNA (**Fig 4C**). The distance between the two COMs of the BRs on the nonspecific DNA (22.7±0.8 Å) remains similarly as that on the specific DNA (22.4±0.4 Å) (**Fig 4D**).

In the specific DNA binding, the Myc BR interacts with both DNA strands along the DNA major groove, with more HBs formed on one strand than on the other (see **Fig 5A**): 9 vs. 2 HBs initially, and 8 vs. 3 later in the simulation. Specific associations with GC13 bases from H359 and E363, respectively, are maintained from the initial stage toward the end; one base association with G11 from R367 initially breaks later.

**Fig 4.**
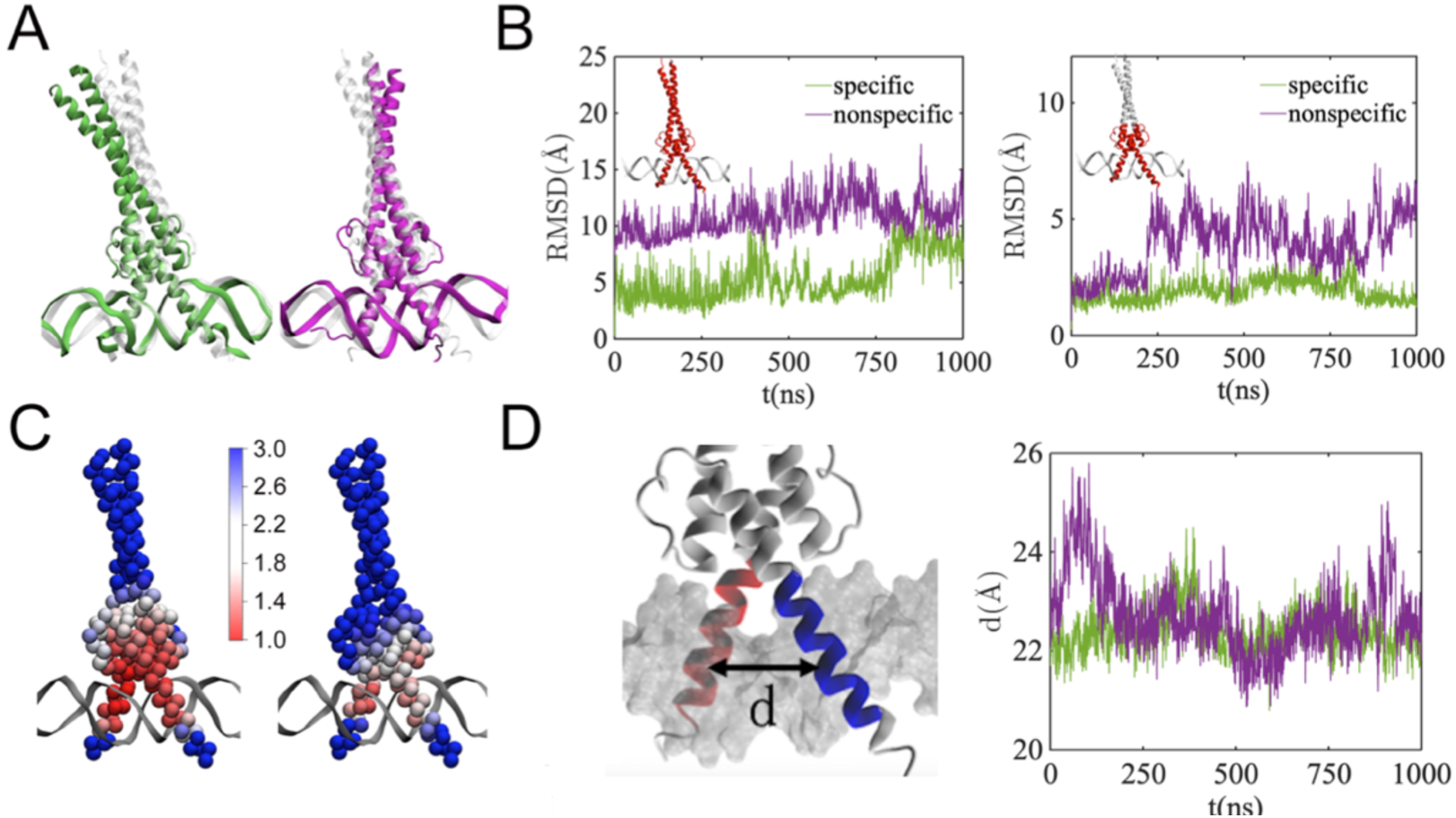
Association of Myc-Max with specific and non-specific DNA sequences from two 1-μs atomistic MD simulations. (A) A comparison of initial (in gray) and final structures of the protein-DNA binding complexes with the specific (left, in green) and non-specific DNA (right, in purple). The initial structures are colored in gray. The structures are aligned between the final and initial ones according to the 10-bp DNA bound nearby the protein. (B) The room-mean-square deviations (RMSDs) of protein. Both the RMSDs of the full protein (left) and the none-Zipper part (right; in red in the inset) are shown, for both specific (green) and non-specific (purple) binding systems. (C) The room-mean-square fluctuations (rmsfs) of the protein bound on the specific (left) and the nonspecific (right) DNA, with colors from red to blue showing the rmsf value from low to high. (D) The distance between the COMs of the two BRs from Myc and Max during the simulations.

**Fig 5.**
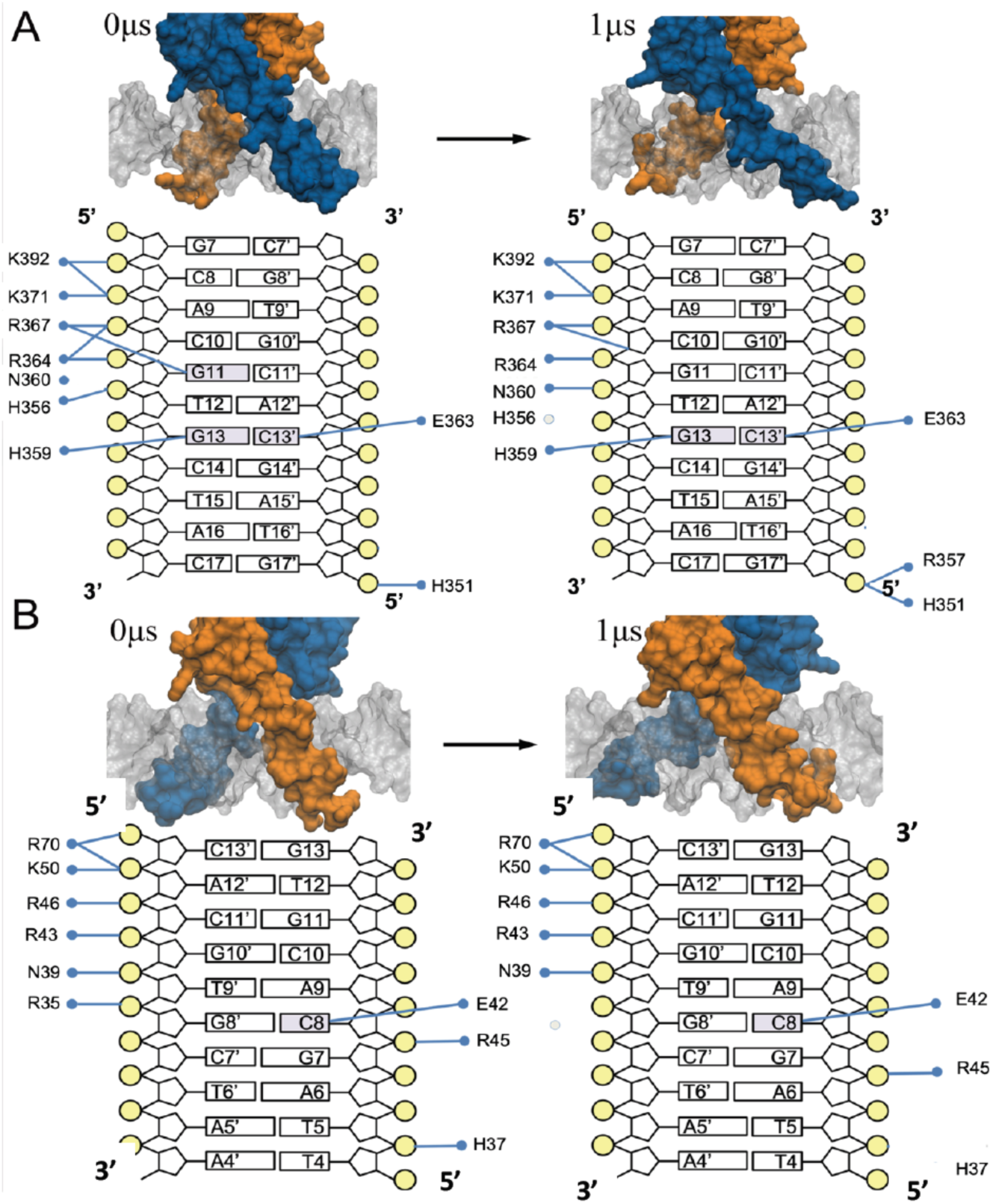
The hydrogen bonds (HB) between Myc-Max and specific (E-box) DNA in the all-atom MD simulation. (A) The HB interactions during the first half (500ns; *left*) and the last half (500ns; *right*) of the simulation between Myc and the DNA. The Myc, Max, and DNA are shown in blue, orange, and gray surfaces, respectively. In the schematics of the HBs at the protein-DNA interface, the DNA bases, sugar, and phosphates are shown by squares, pentagons, and circles, respectively. The DNA bases in contact with protein are colored gray. The HB interaction is defined by a cut-off distance of 3.5 Å between the donor and acceptor atoms and an associated donor-hydrogen-acceptor angle of 140° (the HBs formed >20% the simulation time are shown). (B) The HB interactions during the first half (*left*) and the last half (*right*) of the simulation between Max and the specific DNA.

In comparison, the Max BR maintains 6-7 HBs with the complementary DNA strand (the less associated strand with Myc), and 2-3 HBs with the other strand in the specific DNA binding, with E42 forming HB specifically with the base of C8 (see **Fig 5B**).

For the Myc BR binding on the nonspecific or poly-AT DNA, the protein chain forms 9 HBs with one strand initially, and 6 HBs remain toward the end of the simulation. There has been no stable HB formed between Myc and the other strand, with no base association either (**Fig 6A**). Interestingly, stepping of Myc on the associating strand is initiated (toward 5’ direction), as most of HBs formed initially between Myc and the DNA strand shift for one nucleotide (see **Fig 6A** *right*). As a result, a significant deviation of Myc from the less associated or complementary DNA strand is found, and later Myc dissociates from that strand. Both the initial stepping and the strand dissociation may facilitate further diffusion of the protein on the DNA, which is not sampled in current atomic simulation. In comparison, Max keeps 5-6 HBs with the complementary poly-AT DNA strand (the less associated one with Myc) and 1 HB with the other, which shifts 2 nt distance (toward 5’ on the Myc associating strand; see **Fig 6B** *right*). Overall, Myc and Max are able to form similar amounts of HBs on the two complementary DNA strands, respectively, binding on non-specific DNA.

**Fig 6.**
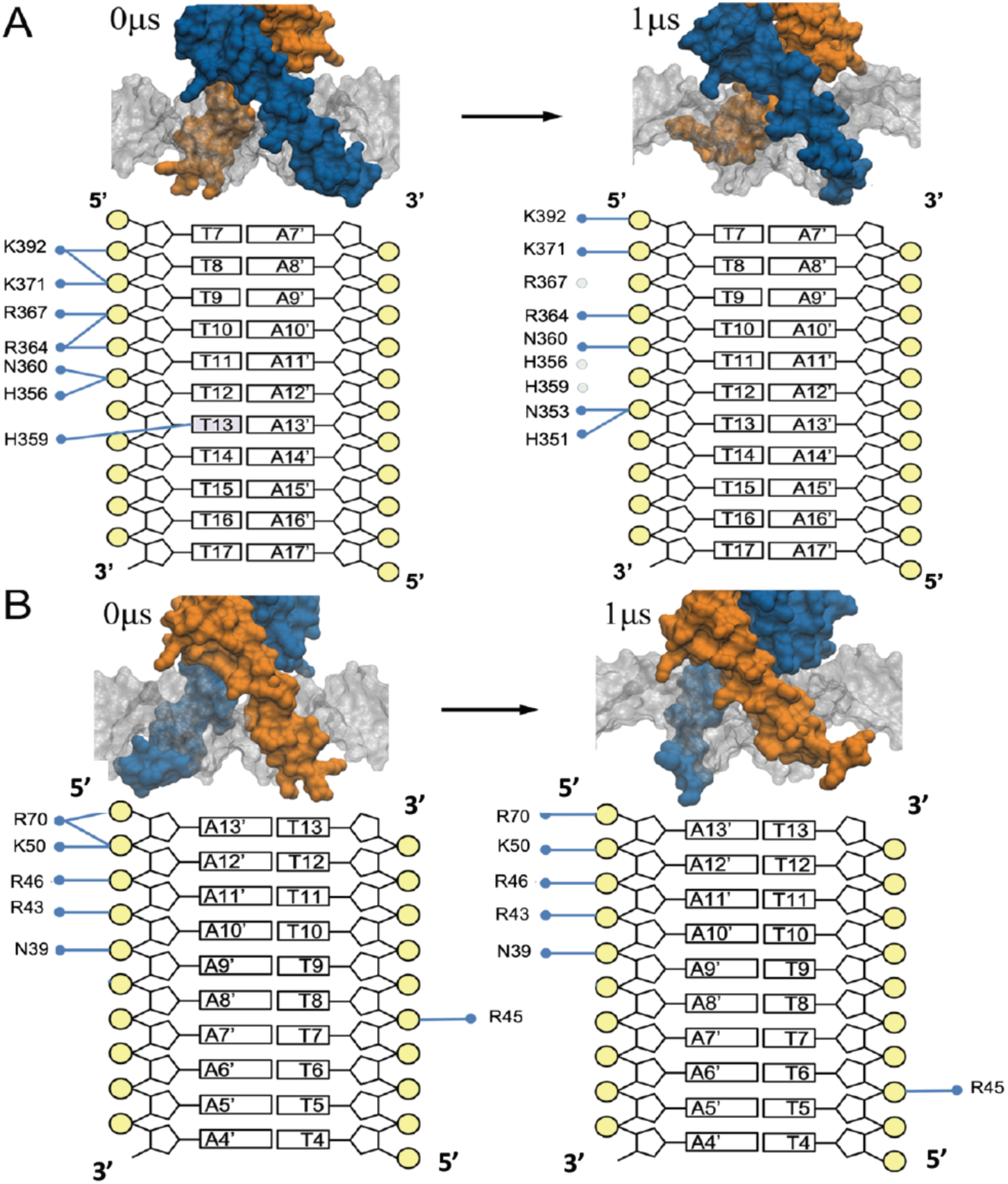
The hydrogen bonds (HB) between Myc-Max and non-specific (poly-AT) DNA in the all-atom MD simulation. (A) The HB interactions during the first half (500ns; *left*) and the last half (500ns; *right*) of the simulation between Myc and the DNA. The coloring, the DNA schematics, and the HB interactions are the same as in Fig 5. (B) The HB interactions during the first half (*left*) and the last half (*right*) of the simulation between Max and the non-specific DNA.

## 3. Discussion

By conducting protein structure-based simulations, we show that diffusion of the Myc-Max heterodimer along DNA can be achieved in inchworm stepping motions. The two BRs of the Myc-Max heterodimer alternate stepping on the DNA, transiting between a closed and an open conformation, i.e., the leading BR steps first (from the closed to open) and the trailing BR follows (open to closed). Such inchworm stepping motions have been identified previously in nucleic acid molecular motors such as PcrA helicase [50, 51] and chromatin remodeler [52]. In PcrA helicase, for example, domain 1A and 2A alternative their affinities with the single-stranded (ss)DNA, in coordination with ATP binding and product release cycles, so that the helicase moves directionally via the inchworm motions along ssDNA [50, 51]. The movements of TF, however, relies essentially on thermal fluctuations, and there is no directional bias for the Myc-Max diffusion along DNA, so that the inchworm motions are sustained in both directions.

Compared with hand-over-hand stepping motions revealed for the cytoskeleton motors myosin and kinesin, which step at about twice the inter-domain distance along the track, the step size in the inchworm protein cannot be larger than the inter-domain distance. Indeed, the step size matches the difference between the inter-domain distances in the open and closed states. Due to substantial electrostatic interactions at protein-DNA interface, the energy barrier for conducting the hand-over-hand type of protein motions can be significantly larger than that for the inchworm motions. Consequently, dimeric TFs and motor proteins moving along nucleic acid track may be more prone to the inchworm stepping motions than to the hand-over-hand motions, no matter in the TF diffusion or the directional movements of the motor protein.

Meanwhile, stochasticity plays a significant role in biomolecular systems. Aside from random directional reversals of the protein during diffusion (i.e. forward to backward), there are occasional BR swapping events that exchanges the left-right positioning of the two BRs. The BR swapping cannot happen, however, directly from the most stabilized closed state. Instead, the closed state needs to transit to a less stabilized tightly-closed state first, as the trailing BR moving forward prior to the leading BR. The BR swapping then happens upon substantial rotation of the heterodimer around its own axis on the DNA, followed by conformational relaxation such that the left-right positioning of the two BRs reverses.

In the heterodimer diffusion, Myc and Max act equally likely to be a leading chain. The left-right positioning of the heterodimer in opposite directions on the dsDNA also appears equivalent. Thus, no obvious bias overall exists between Myc and Max as the heterodimer diffuses along DNA. In the atomic simulation of Myc-Max binding with non-specific DNA, we have noticed that the Myc BR starts stepping on one strand of DNA while dissociating from the other strand. The HB interactions between Myc and its association DNA strand and that between Max and the complementary DNA strand are comparable. That is, no particular bias between two BRs in association with DNA arises even when detailed protein-DNA interactions are considered. Accordingly, inchworm stepping motions revealed in the CG simulations are expected to be maintained similarly in more realistic conditions.

To maintain such inchworm motions, the ‘closed-open-closed’ conformational changes are essential, which highlight the closed and open conformations being crucial for the Myc-Max heterodimer function. The highly populated closed state presented in our CG simulations corresponds well to that being captured in the crystal structure of the Myc-Max DNA binding complex. In such a structure, two BRs bind to the two sides of a same major DNA groove, and the distance between the two BRs (∼ 20 Å) is slightly larger than one half DNA helical pitch. The open conformation, however, has not been captured yet for Myc-Max, though it frequently shows in current CG simulations. In such an open state, the two BRs bind into the neighboring major and minor grooves on the DNA, respectively, with a distance in between (∼33 Å) almost the length of a full DNA helical pitch. In a recent NMR integrated structural dynamics study which measured the Max-Max homodimer conformational fluctuations in association with DNA, it is found that the opening of the binding cleft varies in width by about 2–5 nm [40]. Such an opening of Max-Max may also apply for Myc-Max. In addition, a transiently populated tightly closed state of Myc-Max also appears in our simulations, as the trailing BR moves further close to the leading BR before the leading one moves to open. Via such a destabilized state, stochastic BR swapping for the heterodimer left-right repositioning happens. Accordingly, one expects that the tightly closed state can be captured via fast dynamics detection in the future.

Although we have currently identified stepping or diffusion dynamics of the Myc-Max heterodimer along DNA, how it is linked to its overall DNA search dynamics across genome is not clear. Studies using single-molecule imaging and tracking assay have found c-Myc a global or non-compact explorer of nucleus [53, 54], which leaves a significant number of available sites unvisited, while a compact explorer performs redundant local space exploration. In the inchworm stepping of Myc-Max we capture along DNA, the two BRs move across the DNA strand instead of tracking in the groove in a rotation-coupled manner. Consequently, Myc-Max steps up to ∼ 4 to 5 bps most of time, which may facilitate comparatively fast protein diffusion along DNA, while some of DNA sequences may not be very closely scanned or scrutinized. Whether such stepping behaviors of Myc-Max along DNA impacts on its search dynamics in nucleus is to be investigated in future studies.

## 4. Methods and materials

### 4.1 Coarse-grained simulations

All the CG simulations were performed by using the CafeMol 3.0 software [25]. The initial structure of the Myc-Max heterodimer was taken from the crystal structure (pdb: 1NKP)[38]. The CG protein structure was constructed by using the Go model [48], with each CG particle located on the Cα atom to represent one amino acid. The 3SPN.1 model was used to construct the 200-bp DNA model, with 3 CG particles corresponding to base, sugar and phosphate of one nucleotide [26]. The excluded volume and electrostatic interactions are considered between different subunits. All the CG simulations are conducted by Langevin dynamics.

### 4.2 Setup of atomistic MD simulations

All-atom MD simulations were performed by using the GROMACS 5.12 software [55] with AMBER99SB_ILDN force field [56]. The initial structure of the Myc-Max DNA binding complex was obtained directly from the crystal structure (pdb: 1NKP) [38].

In building of an initial model of Myc-Max binding on the specific DNA, we extended the DNA in the crystal structure to 34 bp to avoid end effects of DNA, by adding random nucleotides to the two ends of the DNA structure in a standard B-form and using the W3DNA web-server (http://w3dna.rutgers.edu/rebuild/regseq) [57]. In building of an initial structure of Myc-Max binding on the nonspecific DNA, we converted the 34 bp DNA above to identical sequences of A and T as poly-AT.

Before MD simulations, the protein-DNA complex was solvated with TIP3P water in a cubic box, and a minimum distance from the complex to the wall was set at 7 Å. We neutralized the systems with Na+ and Cl^−^ to an ionic concentration of 0.15 M. The total system consists of 111,548 atoms for Myc-Max on the specific E-box sequence, and 106,998 atoms for it on the nonspecific poly-AT DNA. The simulations were conducted under the periodic boundary condition. The van der Waals (vdW) and short-range electrostatic interactions used a cutoff distance of 12 Å. The particle-mesh Ewald (PME) method was applied to deal with the long-range electrostatic interactions [58]. The solvated system was minimized with the steepest-descent algorithm, followed by 10-ps MD simulation under the microcanonical or NVE ensemble with a time step 1 fs. After that, a 5-ns equilibrium simulation under the canonical or NVT ensemble and another 5-ns equilibrium simulation under the NPT ensemble with a time step of 1 fs were performed, and position restraints on the heavy atoms of protein and DNA were imposed during the simulation. In the constant temperature simulations, the temperature was set to 300 K using the Langevin dynamics method [59]. The pressure was set at 1 bar using the Nosé-Hoover Langevin piston pressure control method [60]. Finally, the unconstrained 1-μs equilibrium simulations were carried out under the NPT ensemble with a time step of 2 fs.

## Acknowledgements

This work has been supported by NSFC Grant #11775016 and #11635002. JY has been supported by the CMCF of UCI via NSF DMS 1763272 and the Simons Foundation grant #594598 and start-up fund from UCI. We acknowledge the computational support from the Special Program for Applied Research on Super Computation of the NSFC Guangdong Joint Fund (the second phase) under Grant No. U1501501 and from the Beijing Computational Science Research Center (CSRC).

